# A map of gene expression in neutrophil-like cell lines

**DOI:** 10.1101/204875

**Authors:** Esther Rincón, Briana Rocha-Gregg, Sean R. Collins

## Abstract

**Background:** Human neutrophils are central players in innate immunity, a major component of inflammatory responses, and a leading model for cell motility and chemotaxis. However, primary neutrophils are remarkably short-lived, limiting their experimental usefulness in the laboratory. Thus, human myeloid cell lines have been established and characterized for their ability to undergo neutrophil-like differentiation *in vitro*. The HL-60 cell line and its PLB-985 sub-line are commonly used as a model for human neutrophil behavior, but how closely gene expression in differentiated cells resembles that of primary neutrophils has remained unclear.

**Results:** In the present study, we compared the effectiveness of differentiation protocols and used RNA sequencing (RNA-seq) to compare the transcriptomes of HL-60 and PLB-985 cells with published data for human and mouse primary neutrophils. Among commonly used differentiation protocols for neutrophil like cell lines, addition of dimethyl sulfoxide (DMSO) gave the best combination of cell viability and expression of markers for differentiation. However, combining DMSO with the serum-free-supplement Nutridoma resulted in an increased chemotactic response and cell surface expression of the neutrophil markers FPR1 and CD11b without a cost in viability. RNA-seq analysis of HL-60 and PLB-985 cells before and after differentiation showed that differentiation broadly increases the similarity in gene expression between the cell lines and primary neutrophils. Furthermore, the gene expression pattern of the differentiated cell lines correlated slightly better with that of human neutrophils than the mouse neutrophil pattern did. Finally, we created a publicly available gene expression database that is searchable by gene name and by protein domain content, where users can compare gene expression in HL-60, PLB-985 and primary human and mouse neutrophils.

**Conclusions:** Our study verifies that a DMSO-based differentiation protocol for HL-60 and PLB-985 cell lines gives superior differentiation and cell viability relative to other common protocols, and indicates that addition of Nutridoma may be preferable for studies of chemotaxis. Our neutrophil gene expression database will be a valuable tool to identify similarities and differences in gene expression between the cell lines and primary neutrophils, to compare expression levels for genes of interest, and to improve the design of tools for genetic perturbations.

## Background

Neutrophils are the most abundant immune cell population circulating in the human bloodstream, representing 50-70% of all leukocytes. They are generated from myeloid precursors in the bone marrow, where they undergo several stages of maturation, namely myeloblast, promyelocyte, myelocyte, metamyelocyte, band cell and finally polymorphonuclear neutrophil [1]. From the bloodstream, neutrophils are rapidly recruited to sites of inflammation, migrating up gradients of chemical cues in a process called chemotaxis. Neutrophils are among the world’s most accurate chemotaxing cells, and serve as a leading model for eukaryotic chemotaxis [2]. Once at the inflammation site, neutrophils respond to the chemoattractant signals with cytotoxic and inflammatory responses, including secretion of granules (degranulation), ingestion of microbes and other particles (phagocytosis), and generation of neutrophil extracellular traps that catch and kill extracellular microbes. Neutrophil responses require a fine balance: underactivity leaves the body vulnerable to invasion, while over-activity is implicated in inflammatory and autoimmune diseases, and cancer progression [1][3].

Despite much interest in understanding the molecular mechanisms underlying human neutrophil behaviors, their use in experimental research is limited by their very short life-span. They begin to undergo apoptosis within 6 to 12 hours after isolation [4], making techniques such as genetic perturbation impractical. To address this issue, human myeloid cell lines have been established and characterized for their ability to undergo neutrophil-like differentiation *in vitro* [5][6]. The HL-60 cell line, established in 1977 from an acute myeloid leukemia patient, is one of the most commonly used [7]. HL-60 cells are at the myeloblast stage of development but can be induced to differentiate terminally *in vitro* to either a neutrophil-like or a monocyte-like state. Neutrophilic differentiation causes a decrease in cell size, increased nuclear pyknosis and segmentation, and alterations in gene and protein expression. This *in vitro* differentiation makes HL-60 cells a good model to study neutrophil behaviors such as chemotaxis and phagocytosis [8][9][10], as well as the myeloid differentiation process itself [11]. A second commonly used cell line, PLB-985, was originally reported as being established in 1985 from a different patient [12]. However, further characterization determined that PLB-985 is actually a sub-line of HL-60 [13]. PLB-985 cells have similar, but slightly different, properties from those of HL-60 cells, and they are used to study processes including neutrophil ROS production [14] and chemotaxis [15][16][17]. These cell lines serve as primary model systems, but even so, they do not fully recapitulate all neutrophil behaviors [18][6]. Important outstanding questions include to what degree the *in vitro* differentiated cells recapitulate neutrophil gene expression, and what differences in gene expression exist that may affect particular neutrophil behaviors.

Additionally, it is still not clear which of the commonly used reagents and protocols best induces differentiation of HL-60 and PLB-985 cells into a neutrophil-like state. Dimethyl sulfoxide (DMSO), all-trans retinoic acid (ATRA), and dibutyryl cyclic adenosine monophosphate (dbcAMP) are all commonly used to induce differentiation [19][12][6]. While ATRA and dbcAMP are thought to regulate transcription through cognate retinoic acid receptors and cAMP signaling, respectively, the mechanism for DMSO is much less clear. DMSO may act by altering membrane permeability and microviscosity, by affecting synthesis of nucleic acid and proteins, or by inducing changes in chromatin [20][21][22]. Additional reagents have also been reported to improve the efficiency of differentiation. In particular, addition of caffeic acid (CA) [23], replacement of serum with the supplement Nutridoma [24], and stimulation with the cytokine granulocyte colony-stimulating factor (G-CSF) [25] have each been used to promote neutrophilic differentiation in some contexts.

We wanted to address the above outstanding questions by first determining an optimal protocol for efficient neutrophilic differentiation, and then characterizing gene expression in differentiated cells for systematic comparison with primary neutrophils. Our study indicates that DMSO-based differentiation achieves the best balance of neutrophil marker expression with high cell viability and function, and that the use of Nutridoma can further increase cell surface expression of at least some neutrophil markers, including the Formyl Peptide Receptor 1 (FPR1). Our RNA sequencing (RNA-seq) analysis of HL-60 and PLB-985 transcriptomes confirmed that differentiation markedly increases the correlation in gene expression between the cell lines and human neutrophils, slightly exceeding the correlation between mouse and human neutrophil gene expression. Finally, we organized our data, together with previously published data, in a public neutrophil gene expression database that is searchable by gene name and by protein domain content. This database will serve as a resource for the community for easy comparison of gene expression in the cell lines versus human and mouse primary neutrophils, and as a reference for improving gene perturbation strategies in neutrophil models.

## Results and discussion

### Comparison of commonly used differentiation protocols

Since neutrophilic differentiation is often associated with increased cell death, we set out to determine which differentiation protocol produces the most highly penetrant differentiation while maintaining high cell viability. We compared the most commonly used differentiation agents in PLB-985 cells by measuring their ability to induce expression of neutrophil surface markers and reduce proliferation while maintaining cell viability. Initially, we compared differentiation for 6 days with DMSO, ATRA (with 0.5% N,N-Dimethyl formamide (DMF)) and dbcAMP [12][26][27].

To assess differentiation, we measured both early and late differentiation markers. We chose CD11b (Integrin Alpha M, ITGAM) as an early marker, since it begins to appear within three days of differentiation with DMSO [28], and FPR1 as a late neutrophil differentiation marker since it is usually only detectable after five or six days of differentiation. We measured both markers by flow cytometry, using a monoclonal antibody against CD11b and a fluorescent peptide ligand for FPR1 (FLPEP). While undifferentiated cells did not express CD11b or FPR1, a substantial percentage of cells expressed both markers when treated with any of the three differentiation inducers (Figure 1a). Treatment with either DMSO or ATRA caused efficient expression of CD11b, while dbcAMP only induced a bimodal population for this early differentiation marker (Figure 1a). FPR1 was induced with partial penetrance by all differentiation agents. In each case, greater than 50% of cells expressed the receptor, but a sizeable fraction of cells with low or undetectable receptor levels also remained (Figure 1b and Table 1).

**Figure 1.**
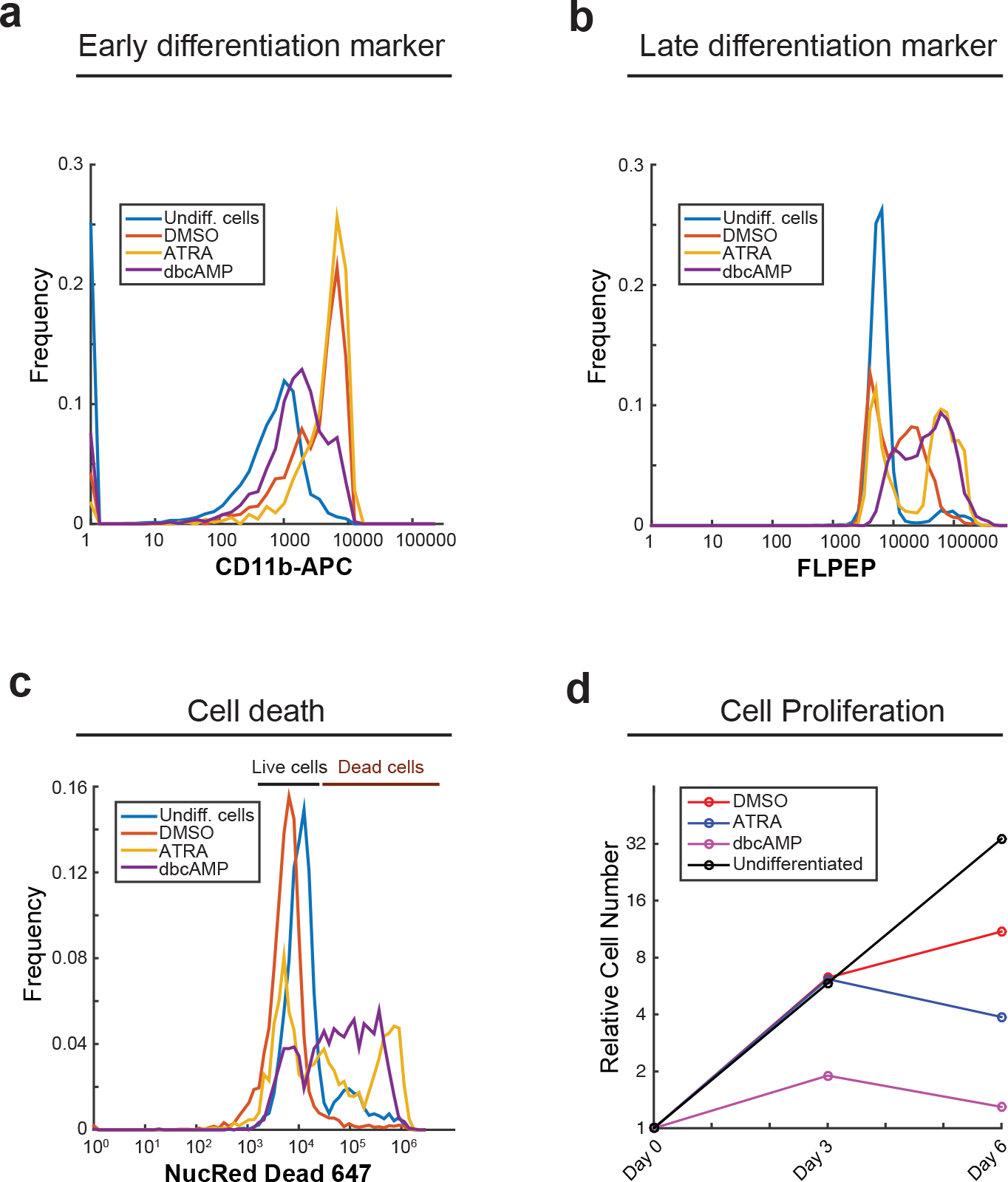
DMSO-based differentiation protocol gives the best differentiation potential based on expression of differentiation markers and cell viability. PLB-985 cells were differentiated into a neutrophil-like state by culturing in different media (DMSO, ATRA or dbcAMP) for 6 days. Undifferentiated cells were also analyzed. Cells were stained with an antibody against CD11b, chosen as an early differentiation marker **(a)**, with FLPEP (a fluorescent ligand of FPR1), chosen as a late differentiation marker **(b)**, or with NucRed Dead 647 Probe, to measure cell death **(c)**. Samples were measured by cytometry and data was analyzed using MatLab. **(d)** Cell growth was monitored at days zero, three, and six of the differentiation protocols, using the trypan blue dye exclusion test. The number of cells were normalized to the initial number of cells. These experiments were performed three times and a representative experiment is shown.

**Table 1.** PLB-985 cell death and expression of FPR1 under several differentiation inducer agents. PLB-985 cells were differentiated into a neutrophil-like state by culturing in the indicated different media for 6 days. Undifferentiated cells were also analyzed. Cells were stained with NucRed Deatd 647 Probe to measure cell death and with FLPEP, a fluorescent ligand of FPR1. Samples were measured by cytometry and data was analyzed using MatLab. For the mean fluorescence intensity (MFI), values were normalized against the MFIof the sample of cells differentiated only with DMSO.

| Differentiation protocol | In NucRed negative population |  | % Cell death (NucRed Dead +) |
| --- | --- | --- | --- |
|  | % Cells expressing FPR1 | Mean Fluorescence Intensity for FPR1 |  |
| Undifferentiated cells | 10 | 33 | 10 |
| DMSO | 52 | 100 | 8 |
| ATRA | 63 | 294 | 44 |
| dbcAMP | 85 | 283 | 58 |
| DMSO + ATRA | 44 | 44 (*) | 70 |
| DMSO + ATRA + CA | 52 | 142 (*) | 73 |
| DMSO + Nutridoma CS | 70 | 720 | 10 |

(\*) Autofluorescence in non-stained samples is very high, probably due to ATRA and CA

We measured the loss of cell viability by staining cells with the NucRed Dead 647 reagent, which binds to DNA in cells with compromised plasma membrane integrity. While DMSO caused little if any apparent cell death, ATRA and dbcAMP each resulted in about 50% inviability in the cell population (Figure 1c and Table 1). We also monitored cell proliferation by counting viable cells using the trypan blue exclusion method. All three treatments reduced proliferation, with dbcAMP acting the most rapidly (Figure 1d). However, by day 6, the number of cells in the ATRA and dbcAMP conditions had markedly decreased, consistent with high rates of cell death. These results also coincide with reports from the initial characterization of the PLB-985 cell line [12].

While our initial results indicated that DMSO is the most suitable agent for inducing neutrophil marker expression without hampering cell viability, we wanted to explore variations of the protocol that might further improve differentiation efficiency. We pursued combinations of DMSO, ATRA [29] and CA [23], as these have been reported to improve differentiation. However, we observed increased cell death and marginal improvement of FPR1 expression at best compared to DMSO alone (Table 1). In an attempt to decrease cell death while still inducing differentiation [30], we limited induction with ATRA to 12h followed by DMSO differentiation media for 5 days. However, the fraction of cells expressing FPR1 was no higher than for DMSO alone (data not shown).

### Replacing serum with Nutridoma during differentiation increases FPR1 surface expression and chemotactic efficiency

We also tested the effects of serum and serum replacement on differentiation. High concentrations of serum have been reported to inhibit differentiation [24]. However, the cell death rate increases markedly under serum-free conditions [24]. Nutridoma, a serum free differentiation supplement, has been reported to improve cell differentiation while providing serum-like support of cell viability [24][31]. To test how serum concentration and Nutridoma affect differentiation in PLB-985 cells, we analyzed upregulation of FPR1 upon differentiation in media containing DMSO, 0.5% fetal bovine serum (FBS), and 2% Nutridoma. Surprisingly, replacing serum with Nutridoma increased the percentage of cells that express FPR1 by approximately 20% compared to DMSO alone (Table 1 and Figure 2a), while cell viability remained similar to that of non-differentiated cells (Table 1). Consistent with our FPR1 results, we observed that the presence of Nutridoma in the differentiation media also increased the surface expression of CD11b (Figure 2b).

**Figure 2.**
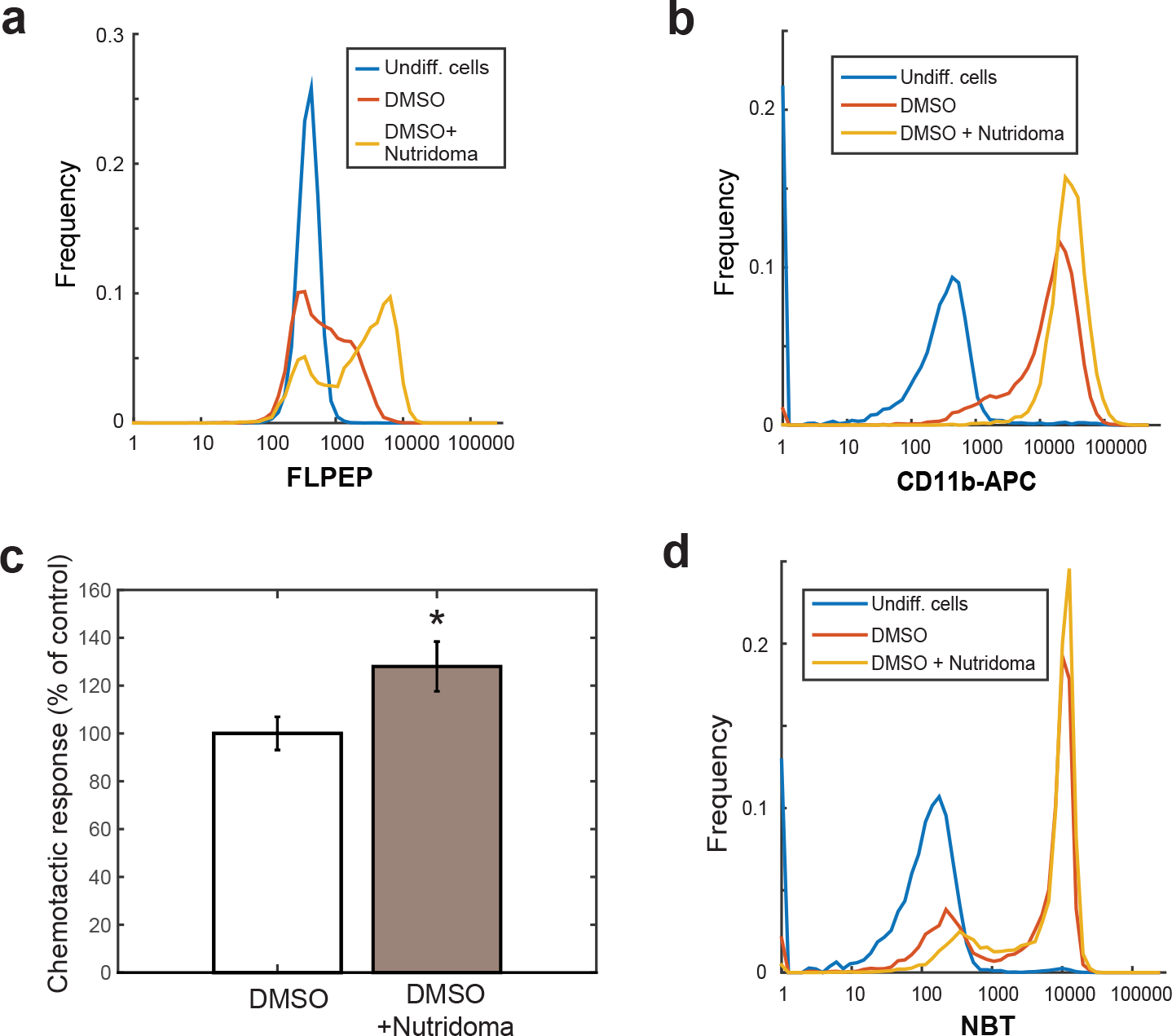
Replacing serum with Nutridoma during differentiation increases FPR1 surface expression and chemotactic efficiency. PLB-985 cells were differentiated into a neutrophil-like state by culturing in media supplemented with 1.3% DMSO and 9% FBS or supplemented with 1.3% DMSO, 2% Nutridoma and 0.5% FBS, for 6 days. Then cells were stained with FLPEP **(a)** or an antibody against CD11b **(b)** and measured by cytometry. Data was analyzed using MatLab. The experiment was repeated three times and a representative experiment is shown. **(c)** Cells differentiated as in (a) and (b) were plated under agarose and analyzed in an automated chemotaxis assay by time-lapse microscopy with chemoattractant uncaging of Nv-fMLF. A cell directionality parameter measuring the angular bias of cell movement towards the gradient source is shown. This experiment was performed three times. Error bars represent the standard error of the mean. Images and statistics were processed using custom MatLab software. **(d)** PLB-985 cells were differentiated into a neutrophil-like as in (a) and (b). Cells were then incubated with NBT solution and 5ug/mL of PMA, at 37°C for 15 minutes, and measured by cytometry.

Next, we wanted to determine if the increase in FPR1-expressing cells is correlated with a more general increase in the functional maturation of the cell population. With this in mind, we tested two major behaviors associated with neutrophilic differentiation: chemotaxis and oxidative burst. We used an automated chemotaxis assay to measure the response of PLB-985 cells to a gradient of fMLF, an FPR1 ligand [26]. When cells were differentiated with DMSO supplemented with Nutridoma and 0.5% FBS, we observed a statistically significant increase (approximately 30%) in the directional accuracy of chemotaxis, validating the efficacy of this differentiation media (Figure 2c). To quantify oxidative burst, we measured the ability of PLB-985 cells to produce reactive oxygen species using the nitroblue tetrazolium (NBT) reduction assay. Differentiation with DMSO alone induced more than 70% of the cells to reduce NBT, meaning that they acquired respiratory-burst activity. When Nutridoma was added to the differentiation media, an additional 15% of the cell population reduced NBT (Figure 2d).

### The presence of Nutridoma, rather than the absence of serum, is responsible for increased FPR1 surface expression and chemotactic efficiency

We wanted to determine whether the increased FPR1 expression was due to the reduced serum levels or a positive effect from the Nutridoma supplement. To test this, we titrated the FBS concentration in differentiation media containing fixed concentrations of DMSO and Nutridoma. We found that most of the increase in FPR1 expression was due to the presence of Nutridoma, and FPR1 expression depended only weakly on the concentration of FBS. However, 0.5% FBS gave a slightly higher population of FPR1 positive cells (Additional file 1a), consistent with previous reports [24].

Although the molecular components of Nutridoma are proprietary, we tested the cytokine G-CSF as a candidate active component. G-CSF is a primary regulator of myeloid cell differentiation in humans, and it has been shown to promote neutrophilic differentiation *in vitro* in some contexts [25][32]. To investigate whether it could be the factor in Nutridoma causing the cells to upregulate FPR1 expression, we differentiated PLB-985 cells using DMSO combined with either Nutridoma or G-CSF [33]. However, based on CD11b and FPR1 expression, G-CSF was not only unable to substitute for Nutridoma, but it actually inhibited DMSO-mediated differentiation (Additional File 1b and 1c).

Based on the above results, we have adopted an optimized differentiation protocol with DMSO in media supplemented with 0.5% FBS and 2% Nutridoma.

### Transcriptional profiles of HL-60 and PLB-985 cells

Having validated an optimal differentiation protocol using DMSO and Nutridoma based on expression of surface markers, we next wanted to compare global gene expression in myeloid cell lines to that of primary neutrophils. While another group had previously compared gene expression in HL-60 cells differentiated with either DMSO or ATRA using microarrays [34], no analysis of HL-60 or PLB-985 cell differentiation had yet been completed with the depth and precision that is now possible with RNA-seq. To this end, we extracted total RNA from the cell lines before and after differentiation (as well as at two intermediate time points for PLB-985 cells) and processed it for RNA-seq (Figure 3a). To account for measurement noise and day-to-day variability, we repeated the RNA-Seq experiment on two independent days separated by more than two months. For comparison, we collated raw datasets for primary human neutrophils [35][36][37], undifferentiated HL-60 cells [38], and primary mouse neutrophils [39][40] from previously published studies [41]. We re-analyzed these data sets along-side our own data using identical parameters and normalization (see Methods for details, and Additional Files 2 and 3 for the full data set).

**Figure 3.**
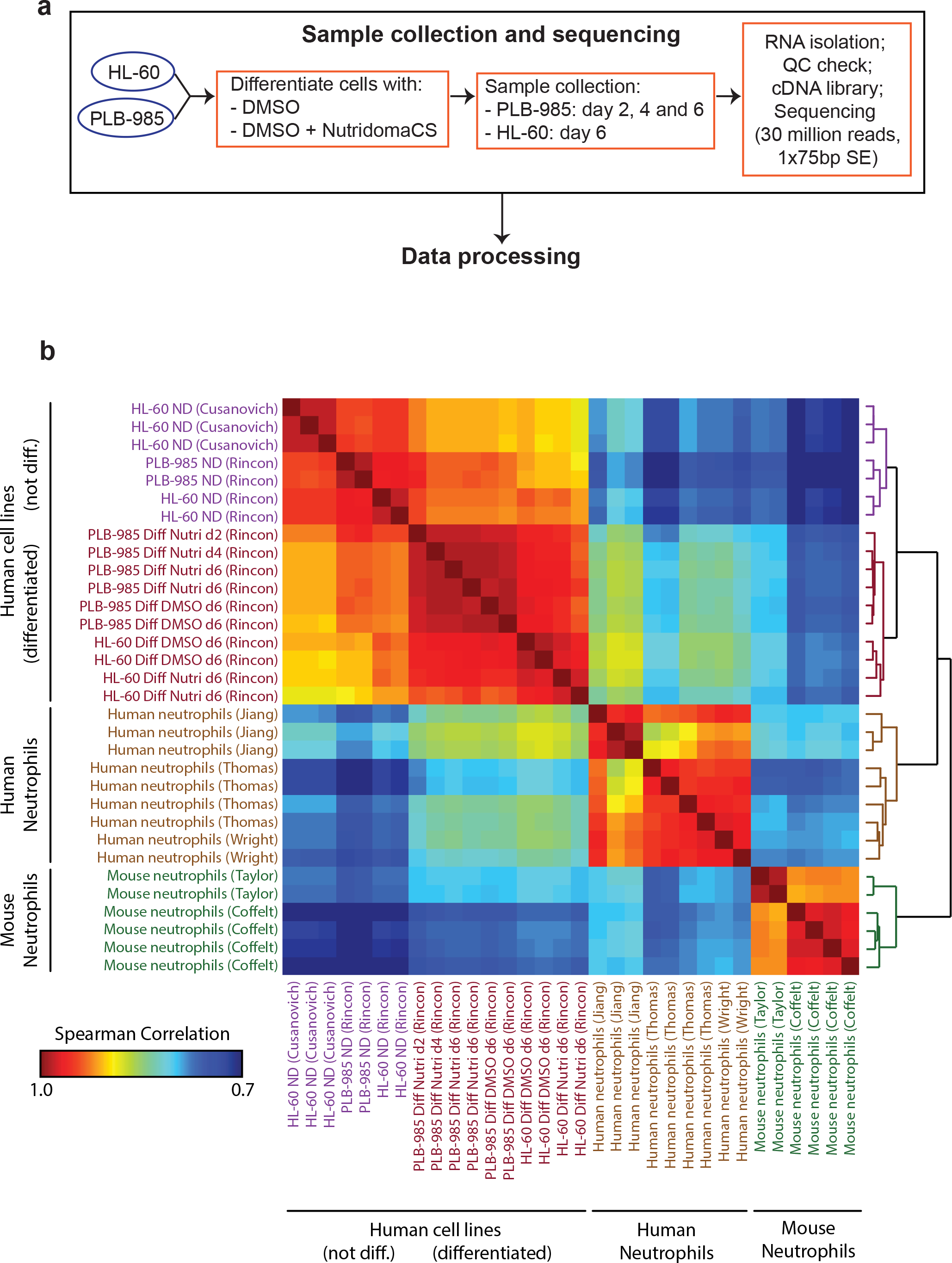
Transcriptional profiles of HL-60 and PLB-985 cells before and after differentiation and comparison to that of primary neutrophils. **(a)** Experimental design: HL-60 and PLB-985 cells were differentiated by culturing in media supplemented with 1.3% DMSO and 9% FBS or 1.3% DMSO, 2% Nutridoma and 0.5% FBS. Total RNA of cells harvested at the indicated time points was isolated. PolyA enrichment and RNA-seq were performed by Applied Biological Materials Inc through their Total RNA Sequencing service. RNA-seq data was subjected to our analysis pipeline as described in Methods. **(b)** Heat map of Spearman correlations between transcriptional profiles of undifferentiated and differentiated HL-60 and PLB-985 cells and human and mouse primary neutrophils, organized by unbiased hierarchical clustering. Color bar indicates correlation strength. Each row/column represents an independent sample.

To compare human and mouse neutrophil transcriptomes, we restricted our analysis to the set of genes mapped as homologs between the two organisms in NCBI’s Homologene database [41]. We then performed unbiased hierarchical clustering of the different transcriptomes using the Spearman rank correlation coefficient as a similarity metric (Figure 3b). As expected, the human primary neutrophil datasets clustered together, as did the mouse primary neutrophil datasets. The cell line datasets also clustered together, with clear subclusters for differentiated and undifferentiated subsets.

### Differentiation results in a neutrophil-like gene expression pattern

The differentiation of the cell lines with either DMSO or DMSO and Nutridoma resulted in large-scale changes in gene expression (Figure 4a), and increased the correlation of their gene expression profiles with that of primary human neutrophils (Figure 3b). The correlation strength increased steadily throughout differentiation, with the biggest change occurring in the first two days (Figures 3b and 4b). To compare the global change in transcription during differentiation, we plotted density-colored scatter plots comparing primary neutrophil data for protein-coding genes to that of undifferentiated or differentiated (with DMSO and Nutridoma) PLB-985 cells (Figure 4c). The plots revealed a global increase in correlation strength, with the bulk of points falling in a tighter more elongated cloud along the diagonal after differentiation.

**Figure 4.**
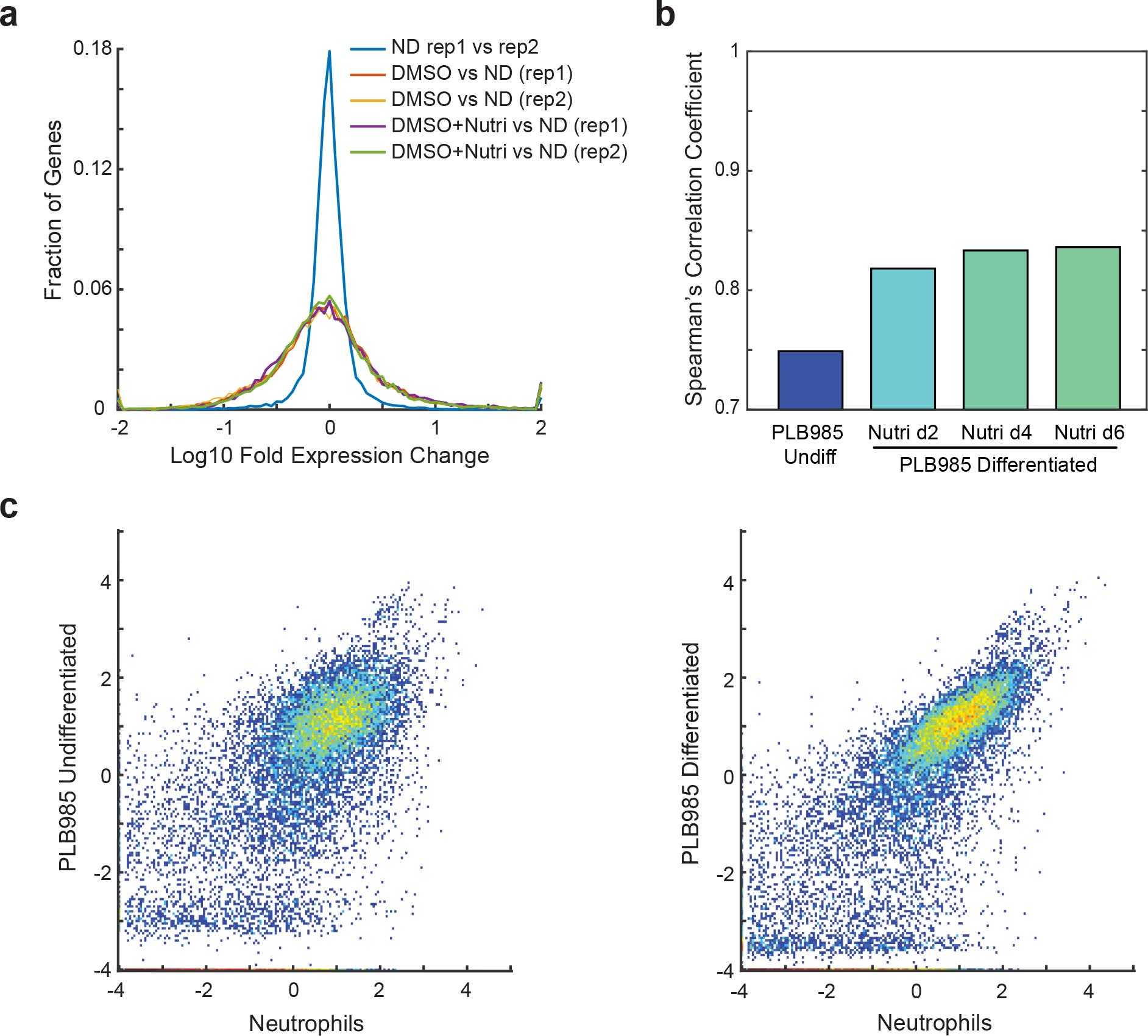
PLB-985 cell differentiation results in neutrophil-like gene expression patterns. **(a)** Histogram of Log10 fold expression changes between replicate measurements of undifferentiated PLB-985 cells (ND), and between undifferentiated cells and those differentiated with either 1.3% DMSO and 9% FBS (DMSO) or DMSO + 2% Nutridoma + 0.5% FBS (Nutri). **(b)** Spearman’s correlation coefficients for the similarity between transcriptional profiles of primary human neutrophils and PLB-985 cells at 0, 2, 4 and 6 days post differentiation with DMSO + 2% Nutridoma + 0.5% FBS. **(c)** Density-colored scatter plot of expression values (normalized FPKM on a log_10_ scale) for protein-coding genes for primary human neutrophils versus undifferentiated PLB-985 cells (left) or those differentiated with DMSO + Nutridoma (right).

As both differentiated human cell lines and primary mouse neutrophils serve as models for human neutrophils, we asked which gene expression pattern correlates more strongly with that of human neutrophils. While both expression patterns correlated more strongly with the human neutrophil data than the undifferentiated cell line data did, we noted that the differentiated cell line data actually correlated more strongly than the mouse neutrophil data (Figure 3b). These results should be taken with some caution, as there may be systematic differences due to different experimental protocols or imperfect mapping of homologs. However, based on global gene expression, our study suggests that the differentiated cell lines resemble human neutrophils as much, or even more so, than mouse neutrophils do. We also noted that while we did see systematic differences between datasets collected by different groups, these differences were generally small relative to the differences between cell types or differentiation states. Indeed, every individual human neutrophil dataset correlated better with the differentiated cell lines than it did with the mouse neutrophil datasets (Figure 3b).

### Nutridoma affects gene expression during differentiation subtly, but consistently in HL-60 and PLB-985 cells

Since the presence of Nutridoma increased cell surface expression of FPR1 and CD11b during differentiation (Table 1, Figure 2a,b), we investigated whether Nutridoma broadly increases neutrophil like differentiation at the level of transcription. First, we compared the expression of FPR1 in our RNA-seq data to surface expression data at different time points during differentiation (Additional file 4a).

Consistent with our earlier observations, the addition of Nutridoma increased the FPR1 surface expression level markedly between day 4 and day 6. However, while Nutridoma also increased FPR1 expression at the mRNA level, this change was considerably smaller than what we observed at the cell surface protein level (Additional file 4a). These results suggest that Nutridoma may influence the cells at the protein level to make them more functionally mature, such as an effect on translation or vesicular trafficking of receptors to the plasma membrane.

Next, we compared global gene expression patterns in cells differentiated with DMSO or with the combination of DMSO and Nutridoma. Although each protocol resulted in gene expression patterns that correlated approximately equally well with the primary human neutrophil data (Figure 3b), we wanted to check if there were reproducible differences in gene expression between the two conditions. To do this, we compared the log10 fold-changes in expression between the two protocols and compared these changes from each of our two replicates of the RNA-Seq experiments. Even though the differences in gene expression were typically modest, they did correlate between the two replicates (Additional file 4b,c). A further analysis of gene annotations using the DAVID functional annotation tool [42], revealed that the set of genes upregulated during differentiation in the presence of Nutridoma (relative to differentiation with DMSO only) are enriched for several annotations including the Gene Ontology (GO) Biological Process annotation of “immune response”, the GO Cellular Compartment annotation “plasma membrane”, and STAT5B transcription factor binding sites (Additional file 5).

Together our observations indicate that, although the correlation in gene expression between primary neutrophils and PLB-985 cells differentiated with or without Nutridoma is roughly equivalent, the presence of this supplement in the differentiation media subtly, but reproducibly affects gene expression at the mRNA level. In addition, Nutridoma also causes non-transcriptional changes which increase the cell surface expression of the neutrophil markers FPR1 and CD11b.

### SNP analysis confirms that PLB-985 is genetically identical to HL-60

Although the PLB-985 cell line was originally described as a distinct cell line established from an acute myeloid leukemia patient in 1985 [12], Drexler and colleagues more recently reported that this cell line is actually a subclone of the HL-60 cell line [13]. The description of PLB-985 as a misidentified cell line can be found in the National Center for Biotechnology Information (NCBI) BioSample database (https://www.ncbi.nlm.nih.gov/biosample/3151776) or in the ICLAC Database of Cross-contaminated or Misidentified Cell Lines (http://iclac.org/databases/cross-contaminations/). The RNA-seq data that we generated for both cell lines allowed us to investigate this reclassification. We used the Genome Analysis Toolkit (GATK) pipeline to call variant genotypes for each sample from our RNA-Seq raw data and from the primary neutrophil data which could serve as a negative control [43]. As expected, we found identical genotypes for replicate samples, and distinct genotypes for primary neutrophil data from different donors. Consistent with previous reports, we also found identical genotypes for the PLB-985 and HL-60 cell lines (Additional File 6). However, even though they are genetically identical, our RNA-Seq data indicates that there are reproducible gene expression differences between PLB-985 and HL-60 cells.

We conclude that PLB-985 is a sub-line of HL-60 with some differences in gene expression which may make it better suited for some experimental investigations.

### A searchable database of neutrophil and neutrophil-like cell line gene expression

A major goal of our study was to create an accessible database of gene expression for neutrophils and neutrophil-like cell lines. Such a tool would allow researchers to assess the expression of gene families of interest, or compare gene expression in cell lines to that in primary neutrophils to verify similarities or identify key differences. To highlight how our compiled datasets can be used, we analyzed expression levels of four sets of genes known to be important for neutrophil functions: cell surface receptors, proteins involved in reactive oxygen production, guanine nucleotide exchange factors (GEFs) for Rho family GTPases, and adhesion receptors (Figure 5a-d). In each case, differentiation broadly increased the similarity in gene expression of these gene sets between the cell lines and primary neutrophils, including upregulation of chemoattractant receptors, oxidative burst-related genes, adhesion molecules, and other neutrophil-associated genes (Figure 5a-d). In contrast, related oxidase components and adhesion receptors not associated with neutrophils remained lowly expressed (Figure 5b,d). However, key differences between the cell lines and primary neutrophils are also apparent. For example, the formyl peptide receptor FPR1 is highly expressed in the cell lines, but expression of the IL-8 cytokine receptor CXCR1 is lacking (Figure 5a). Similarly, many GEFs including PREX1, ARHGEF2 (also known as GEF-H1), VAV1, and ARHGEF6 (also known as alpha-PIX) are expressed at levels close to or equal to that found in neutrophils. However, DOCK5 and PLEKHG3 are expressed at much lower levels.

**Figure 5.**
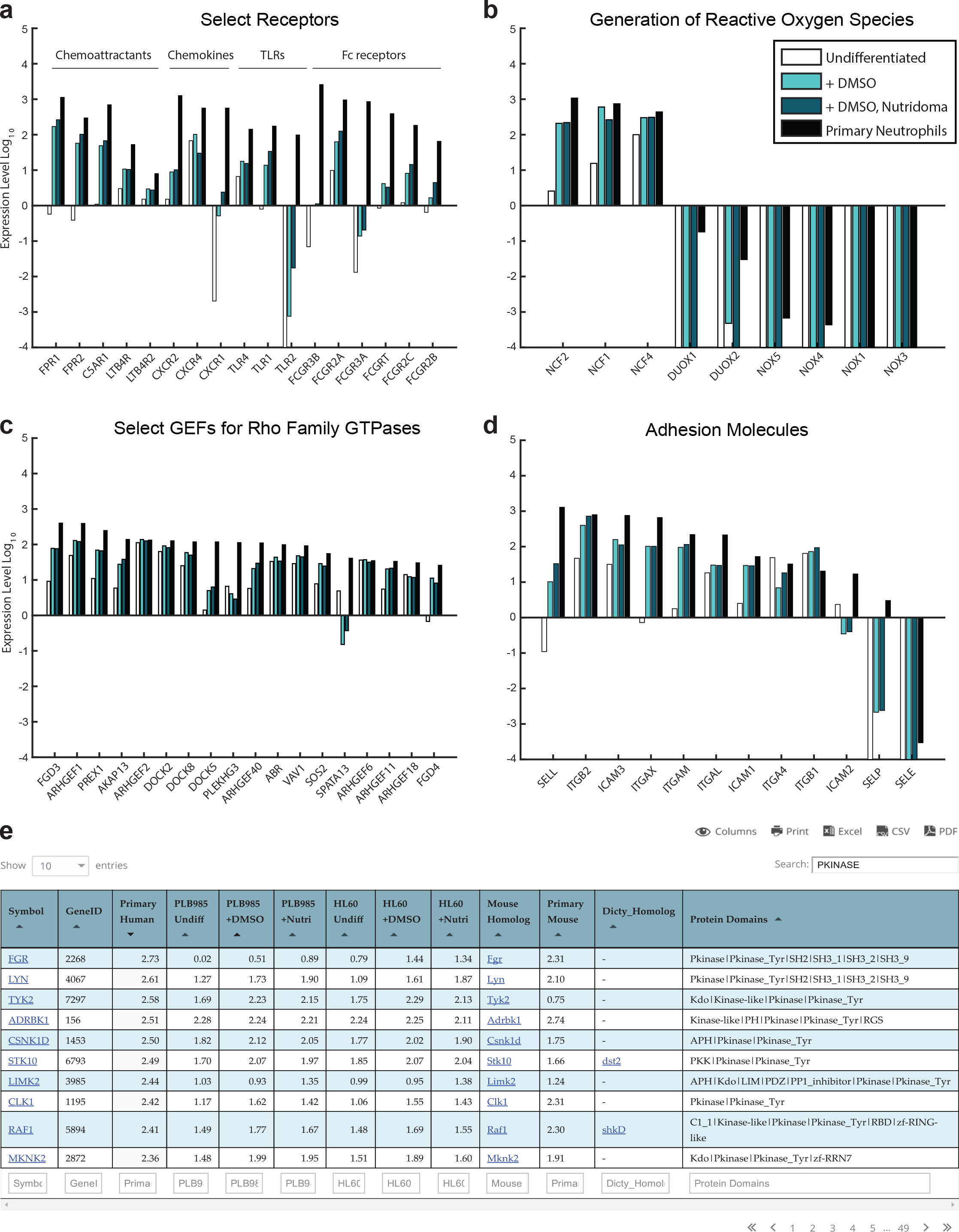
An easily searchable online database of neutrophil and neutrophil-like cell gene expression. **(a-d)**. Gene expression data is shown (normalized FPKM data on a log10 scale) for the indicated genes in undifferentiated PLB-985 cells, PLB-985 cells differentiated with the DMSO only protocol, PLB-985 cells differentiated with the Nutridoma + DMSO protocol, and primary human neutrophils (collated from 3 published studies). Shown is data for select receptors **(a)**, genes related to the production of reactive oxygen species **(b)**, select guanine nucleotide exchange factors (GEFs) for Rho Family GTPases **(c)** and adhesion molecules **(d)**. **(e)** Our collated RNA-seq data is available in an easily searchable form on our lab website: http://collinslab.ucdavis.edu/neutrophilgeneexpression/. Data can be searched by gene name, or by the presence of specific PFAM domains of interest. Data can also be sorted by expression level in any of the samples. In this example, we identify the top 10 genes containing PKINASE domain, ordered by expression level in primary human neutrophils.

Finally, to make our RNA-seq results easily accessible to other researchers, we published a comprehensive, interactive database on our website (http://collinslab.ucdavis.edu/neutrophilgeneexpression/) (Figure 5e). This database is searchable by gene id, symbol, or protein domain and can be sorted by expression level in any of the cell types. In addition to our own RNA-seq data, we included published RNA-seq data for primary human and *Mus musculus* (mouse) neutrophils, and links to mouse and *Dictyostelium discoideum* homologs. *D. discoideum* is an amoeboid protozoan and a valuable model organism for the study of chemotaxis and phagocytosis. This complete merged data set is also available as a table in Additional File 7.

## Conclusions

We first compared established differentiation protocols for myeloid cell lines, with a focus on maximizing cell surface expression of neutrophil markers without compromising cell viability. Our study verifies that a DMSO-based differentiation protocol for HL-60 and PLB-985 cell lines gives superior differentiation and cell viability relative to other common protocols, and indicates that addition of Nutridoma may be preferable for studies of chemotaxis. Our RNA-seq data allowed us to generate a map of gene expression in differentiated HL-60 and PLB-985 cells which could be combined with primary human and mouse neutrophil data from the literature. We annotated this map, creating a database that is searchable by gene name and by protein domain content to allow rapid interrogation of gene expression in neutrophils and neutrophil-like cell lines. Our neutrophil gene expression database will be a valuable tool to identify similarities and differences in gene expression between the cell lines and primary neutrophils, to compare expression levels for genes of interest, and to improve the design of tools for genetic perturbations.

## Methods

### Reagents and antibodies

DMSO, ATRA, CA, dbcAMP, PMA and NBT were purchased from Sigma, Nutridoma-CS from Roche, Trypan Blue solution from Corning, G-CSF from Applied Biological Materials (abm), FLPEP (catalog # F1314) and NucRed Dead 647 from Life Technologies, and Anti-CD11b-APC from Biolegend (Clone ICRF44, catalog # 301309).

### Cell culture and differentiation

HL-60 and PLB-985 cells were obtained as a gift from the laboratory of Dr. Orion Weiner. Generation of the stable cell lines expressing fusions of histone H2B to mTurquoise or mCherry were described elsewhere [26]. The PLB-985 cell line expressing H2B-mCherry was used for all the experiments described throughout the manuscript. Cells were cultured in complete media [RPMI 1640 with HEPES and GlutaMAX (Gibco, catalog # 72400-047) supplemented with 9% fetal bovine serum (FBS) as well as streptomycin (100 mg/mL) and penicillin (100 U/mL) (P/S)] at 37°C in a humidified atmosphere with 5% CO2. Cell cultures were passaged two to three times per week, maintaining cell densities between 10^5^ and 2x10^6^ cells per ml. The cells were differentiated into a neutrophil-like state by culturing at an initial density of 2 × 10^5^ using one of seven media recipes: a) RPMI 1640 complete media supplemented with 1.3% DMSO for 6 days; b) RPMI 1640 media supplemented with 5% FBS, 0.5% DMF and 2 uM ATRA (renewed on day 3 of the 6-day differentiation period); c) RPMI 1640 complete media supplemented with 750 uM dbcAMP for 6 days; d) RPMI 1640 complete media supplemented 1.3% DMSO and 2 uM ATRA for 6 days; e) RPMI 1640 complete media supplemented with 1.3% DMSO, 2 uM ATRA and 52 uM CA for 6 days; f) RPMI 1640 media supplemented with different FBS concentrations (0.5, 2, 5 or 10%), 1.3% DMSO and 2% Nutridoma for 6 days; g) RPMI 1640 media supplemented with 0.5% FBS, 1.3% DMSO and 30 ng/ml G-CSF for 6 days.

### Immunolabeling and cytometry

Differentiation of PLB-985 cells into a neutrophil-like state was assessed by measuring the levels of CD11b and the formyl peptide receptor FPR1 on the cell surface. For detection of CD11b, cells were harvested, washed with phosphate-buffered saline (PBS) and stained with primary antibody antiCD11b-APC (clone ICRF44) for 45 min at 4°C in FACS buffer (0.5% bovine serum albumin and 0.05% sodium azide in PBS). After fluorescent labeling, the samples were washed and acquired with the BD FACSCanto II flow cytometer (BD Biosciences, Franklin Lakes, NJ). Relative levels of FPR1 were assessed by binding of N-formyl-norleucyl-leucyl-phenylalanyl-norleucyl-tyrosyl-lysine-fluorescein (FLPEP) (Life Technologies product number F1314), a fluorescent ligand of the receptor. Cells were mixed with ice cold media containing FLPEP to give a final concentration of 10 nM, placed on ice for 10 min, and acquired with the BD FACSCanto II flow cytometer. For two conditions (DMSO+ATRA and DMSO+ATRA+CA), cellular autofluorescence was high in the fluorescein channel, and it was corrected by subtracting the mean autofluorescence from the unstained control samples. We measured cell death using NucRed Dead 647 Probe (Life Technologies, product number R37113) according to the manufacturer’s protocol. FACS data was analyzed using MatLab (Mathworks, Inc.). To determine the percentage of cells expressing the target, we set thresholds using an appropriate negative control (isotype immoglobulins) for CD11b or non-stained cells for FLPEP and NucRed Dead 647.

### Cell growth

To measure cell proliferation, cells were counted at day three and six of the differentiation protocols, using the trypan blue dye exclusion test. The number of cells were normalized against the initial number of cells.

### Chemotaxis assay

The chemotaxis assay was performed as previously described [26]. Briefly, differentiated PLB-985 cells were plated in 96-well plates under low melting temperature agarose and imaged by time-lapse microscopy with chemoattractant uncaging. A caged UV sensitive derivative N-nitroveratryl derivative (Nv-fMLF) of N-formyl-methioninealanine-phenylalanine was used, and UV photo-uncaging was performed to generate chemical gradients. Image processing and statistical analyses of chemotaxis were performed using custom MatLab software.

### NBT reduction assay

Undifferentiated and differentiated cells were harvested and resuspended in Leibovitz’s L-15 media (ThermoFisher, product number 11415-064) supplemented with FBS. Cells were then mixed with ice cold media containing 2X NBT solution (Sigma, product number N5514) and 5ug/ml of phorbol 12-myristate 13- acetate (PMA) in L-15 media and incubated at 37°C for 15 minutes. After incubation, deposition of purple formazan granules (from the reduction of NBT) was acquired with the BD FACSCanto II flow cytometer.

### RNA-seq analysis pipeline

Total RNA was isolated using the RNeasy Plus Mini Kit (Qiagen) according to the manufacturer’s protocol. PolyA enrichment and RNA-seq were performed by Applied Biological Materials Inc through their Total RNA Sequencing service. The samples were subjected to polyA enrichment followed by fragmentation, first and second strand synthesis, adenylation of 3’ ends, adapter ligation, DNA fragment enrichment, and real-time PCR quantification. For each sample, over thirty million 1×75-base single-end reads were produced on the Illumina NextSeq 500 platform. Human raw RNA-seq reads were aligned to the GRCh38.p5 version of the human genome using the GENCODE GRCh38 (release 24) genome annotations and the STAR two-pass algorithm (STAR version 2.4) [44]. Mouse RNA-seq reads were aligned to the GRCm38 version of the mouse genome using the GENCODE M8 annotations. Gene expression was quantified using the open-source software Cufflinks (version 2.2.1) [45]. Gene expression was normalized using the “uniform genes” strategy of Glusman et al [46]. Briefly, a set of “uniform genes” was defined as all genes with expression levels between the 50^th^ and 90^th^ percentile in all samples. A scaling factor was computed for each dataset as the geometric mean of the expression levels of these “uniform genes.” Each dataset was then normalized by dividing by its own scaling factor and multiplying by the mean scaling factor of the human datasets. Normalization, comparison and correlation of the different transcriptomes, and hierarchical clustering of the data, was done using MatLab. Log base 10 expression values were calculated after normalization, and genes with zero read counts were assigned a value of -4 to avoid infinite values. Hierarchical clustering was done using one minus the Spearman rank correlation coefficient as the distance metric and average linkage. Pairwise Spearman rank correlation coefficients were displayed in the cluster heatmap.

For our final data tables, we averaged the data over replicate samples using the geometric mean. For the primary human and mouse neutrophil datasets, we first averaged the replicate measurement performed by each laboratory group (note that the Thomas et al [37] and Wright et al [36] datasets are from the same laboratory), and then averaged over the different laboratories to give equal weight to each lab to account for potential lab-to-lab systematic differences.

For the density-colored scatter plots in Figure 4a, the averaged gene expression profiles were used. Only protein-coding genes were included in the plot, based on NCBI’s “Gene type” designations (18,976 genes) [41]. Data for noncoding RNAs were much more variable, perhaps in part due to the polyA enrichment step of our experimental processing.

For the cluster of gene expression profiles in Figure 3b, only genes mapped between human and mouse as homologs in NCBI’s Homologene database [41] were included.

Variant calling for genotyping was performed using the GATK pipeline and the GATK best practices guidelines [43]. After variant calling, the genotype data was filtered to keep only loci with a read depth of at least 10 in all samples and a quality score of greater than 100. Genotypes were then compared to determine the fraction of loci with identical genotypes for each pair of samples.

### Generation of the database

Normalized RNA-seq data (prepared as described above) was compiled into a database using MySQL and published on our website (collinslab.ucdavis.edu). *Dictyostelium discoidum* homologs were downloaded from InParanoid (http://inparanoid.sbc.su.se/cgi-bin/index.cgi).

### Accession numbers

HL-60 and PLB-985 undifferentiated and differentiated RNA-seq data were deposited at the NCBI Gene Expression Omnibus (GEO) database (Accession: GSE103706). Previously published data for primary human neutrophils from the Wright et al [GEO: GSE40548], Jiang et al [GEO: GSE66895], and Thomas et al [GEO: GSE70068] studies, for undifferentiated HL-60 cells from the Cusanovich et al study [GEO: GSE68103], and for primary mouse neutrophils from the Taylor et al [GEO: GSE55090] and Coffelt et al [GSE55633] studies were downloaded from NCBI’s GEO database.

## Abbreviations

(RNA-seq): RNA sequencing
(DMSO): Dimethyl sulfoxide
(ATRA): All-trans retinoic acid
(dbcAMP): Dibutyryl cyclic adenosine monophosphate
(fMLF): N-formyl-methionyl-leucyl phenylalanine
(DMF): N,N-Dimethyl formamide
(FPR1): Formyl Peptide Receptor 1
(FLPEP): Fluorescent N-formyl-methionyl-leucyl phenylalanine (fMLF) peptide ligand
(CD11b): Integrin Alpha M, ITGAM
(CA): Caffeic acid
(MFI): Mean fluorescence intensity
(FBS): Fetal bovine serum
(NBT): Nitro Blue tetrazolium
(G-CSF): Granulocyte colony-stimulating factor
(GEF): Guanine nucleotide exchange factor
(SNPs): Single-nucleotide polymorphisms
(GATK): Genome Analysis Toolkit
(NCBI): National Center for Biotechnology Information
(FBS): Fetal bovine serum
(P/S): Streptomycin and penicillin
(PBS): Phosphate-buffered saline
(FACS): Fluorescence-activated cell sorting
(PMA): Phorbol 12-myristate 13-acetate

## Declarations

### Ethics approval and consent to participate

Not applicable

### Consent for publication

Not applicable

### Availability of data and materials

The datasets generated and analyzed during the current study are available in the NCBI GEO database under the accession number GSE103706.

### Authors’ contributions

ER participated in the experimental work, data compilation, analysis and writing of the manuscript. BRG participated in data analysis, generation of the database, and writing of the manuscript. SRC participated in the data compilation, analysis and writing of the manuscript.

## Acknowledgements

We are grateful to colleagues who generously provided reagents. We thank George Bell, Dean Natwick and Michael Pargett for stimulating discussion and critical assessment of the manuscript.

## Additional files

**Additional File 1. (a)** PLB-985 cells were differentiated into a neutrophil-like state by culturing in media supplemented with 1.3% DMSO, 2% Nutridoma and the indicated percentage of FBS. Cells were stained with FLPEP and measured by cytometry. **(b-c)** PLB-985 cells were differentiated into a neutrophil-like state by culturing in media supplemented with 1.3% DMSO 0.5% FBS and either 2% Nutridoma or 30ng/mL G-CSF for 6 days. Cells were stained with anti-CD11b **(b)** or with FLPEP **(c)** and measured by cytometry. Cytometry data was analyzed using MatLab in all the experiments. Data for the undifferentiated cells and those differentiated with DMSO or DMSO + Nutridoma is the same as that shown in Figure 2.

**Additional File 2. Full tables of processed gene expression values measured by RNA-Seq in this study**. This Excel file contains two sheets. The first sheet contains FPKM gene expression values generated by Cufflinks for all samples analyzed by RNA-Seq in this study. The second sheet contains the log10-transformed normalized expression values. These values were computed from the FPKM values as described in the Methods section.

**Additional File 3. Full tables of reanalyzed gene expression data for primary neutrophils and HL-60 cells from previously published studies**. This Excel file contains four sheets. The first sheet contains FPKM gene expression values generated by Cufflinks for all primary human neutrophil and HL-60 samples reanalyzed in this study. The second sheet contains the corresponding log10-transformed normalized expression values. The third sheet contains FPKM gene expression values for all primary mouse neutrophil samples reanalyzed in this study, and the fourth sheet contains the corresponding log10-transformed normalized values.

**Additional File 4. Nutridoma affects gene expression during differentiation subtly, but consistently in HL-60 and PLB-985 cells. (a)** PLB-985 cells were differentiated into a neutrophil-like state by culturing in media supplemented with 1.3% DMSO, 2% Nutridoma and 0.5% FBS. At the indicated days post differentiation, cells were harvested and processed for RNA-seq analysis or stained with FLPEP. Shown are the expression level of FPR1 at the mRNA level (from our RNA-seq data) and at the surface of the cells (FLPEP staining). Values for undifferentiated cells and for cell differentiated with 1.3% DMSO and 9% FBS are shown for comparison. Values were normalized by the values for the sample differentiated with only DMSO. **(b)** Density-colored scatter plots of expression values (normalized FPKM on a log10 scale) for protein-coding genes for HL-60 (left) or PLB-985 cells (right) cells differentiated with DMSO + Nutridoma versus those differentiated with DMSO. These data represent the averages of two replicate experiments. **(c)** Density-colored scatter plots of log10 fold-changes between the two protocols (subtracting DMSO values from DMSO + Nutridoma) comparing each of our two replicates of the RNA-Seq experiments for HL-60 (left) or PLB-985 (right) cells. To avoid noise from lowly expressed genes, only protein-coding genes with a mean log10 normalized expression value of -1 or higher were included. The Pearson’s (R) and Spearman’s correlation coefficients are shown.

**Additional File 5. Summary of enriched gene annotations for the set of genes upregulated in the presence of Nutridoma**. This Excel file contains two sheets. The first contains a set of genes determined to be upregulated in differentiation with Nutridoma relative to differentiation with DMSO alone. This gene set was selected based on a mean log10 normalized expression value of at least -1 across our differentiation samples, and a minimum 3-fold upregulation in each Nutridoma differentiation sample compared to its corresponding DMSO differentiation sample (for both PLB-985 and HL-60 cells). The table contains NCBI GeneIDs, Ensembl GeneIDs, gene names, and the mean log10 normalized expression values in Nutridoma and DMSO differentiation samples. Sheet 2 contains a summary of annotations enriched in for this set of upregulated genes. Annotation enrichment was calculated using the DAVID online gene annotation analysis tool [42].

**Additional File 6. SNP analysis confirms that PLB-985 is genetically identical to HL-60**. Variant calling for genotyping was performed using the GATK pipeline and the GATK best practices guidelines for each sample from our RNA-Seq raw data and from the primary human neutrophil RNA-Seq data. The heatmap indicates the fraction of loci with identical genotypes for each pair of samples (see Methods for more details).

**Additional File 7. Table of cell line and primary neutrophil gene expression data**. This comma separated value file contains the final averaged log10 normalized gene expression values for undifferentiated and differentiated cell lines, as well as primary human and mouse neutrophils. This file contains all of the data included in our online searchable neutrophil gene expression database.

## References

1. Kolaczkowska E, Kubes P. Neutrophil recruitment and function in health and inflammation. Nat. Rev. Immunol. [Internet]. 2013;13:159–75. Available from: http://www.ncbi.nlm.nih.gov/pubmed/23435331

2. Rappel WJ, Loomis WF. Eukaryotic chemotaxis. Wiley Interdiscip. Rev. Syst. Biol. Med. 2009;1:141–9.

3. Mantovani A, Cassatella MA, Costantini C, Jaillon S. Neutrophils in the activation and regulation of innate and adaptive immunity. Nat. Rev. Immunol. 2011;11:519–31.

4. Mollinedo F, Santos-Beneit AM, Gajate C. The Human Leukemia Cell Line HL-60 as a Cell Culture Model To Study Neutrophil Functions and Inflammatory Cell Responses. In: Clynes M, editor. Anim. Cell Cult. Tech. [Internet]. Berlin, Heidelberg: Springer; 1998. p. 264–97. Available from: https://doi.org/10.1007/978-3-642-80412-0_16

5. Collins SJ, Gallo RC, Gallagher RE. Continuous growth and differentiation of human myeloid leukaemic cells in suspension culture. Nature. 1977;270:347–9.

6. Gupta D, Shah HP, Malu K, Berliner N, Gaines P, Gupta D, et al. Differentiation and Characterization of Myeloid Cells. Curr. Protoc. Immunol. [Internet]. Hoboken, NJ, USA: John Wiley & Sons, Inc.; 2014. p. 22F.5.1–22F.5.28. Available from: http://doi.wiley.com/10.1002/0471142735.im22f05s104

7. Collins SJ, Ruscetti FW, Gallagher RE, Gallo RC. Terminal differentiation of human promyelocytic leukemia cells induced by dimethyl sulfoxide and other polar compounds. Proc. Natl. Acad. Sci. U. S. A. 1978;75:2458–62.

8. Hauert AB, Martinelli S, Marone C, Niggli V. Differentiated HL-60 cells are a valid model system for the analysis of human neutrophil migration and chemotaxis. Int. J. Biochem. Cell Biol. [Internet]. 2002;34:838–54. Available from: http://linkinghub.elsevier.com/retrieve/pii/S1357272502000109

9. Fleck RA, Romero-Steiner S, Nahm MH. Use of HL-60 Cell Line To Measure Opsonic Capacity of Pneumococcal Antibodies. Clin. Vaccine Immunol. 2005;12:19–27.

10. Skoge M, Wong E, Hamza B, Bae A, Martel J, Kataria R, et al. A worldwide competition to compare the speed and chemotactic accuracy of neutrophil-like cells. PLoS One. 2016;11:1–19.

11. Birnie GD. The HL60 cell line: a model system for studying human myeloid cell differentiation. Br. J. Cancer. Suppl. 1988;9:41–5.

12. Tucker KA, Lilly MB, Heck L, Rado TA. Characterization of a new human diploid myeloid leukemia cell line (PLB-985) with granulocytic and monocytic differentiating capacity. Blood [Internet]. 1987;70:372–8. Available from: http://www.ncbi.nlm.nih.gov/pubmed/3475136

13. Drexler HG, Dirks WG, Matsuo Y, MacLeod RAF. False leukemia-lymphoma cell lines: an update on over 500 cell lines. Leukemia. 2003;17:416–26.

14. Ashkenazi A, Marks RS. Luminol-dependent chemiluminescence of human phagocyte cell lines: Comparison between DMSO differentiated PLB 985 and HL 60 cells. Luminescence. 2009;24:171–7.

15. Yang HW, Collins SR, Meyer T. Locally excitable Cdc42 signals steer cells during chemotaxis. Nat. Cell Biol. [Internet]. 2015;18:191–201. Available from: http://www.ncbi.nlm.nih.gov/pubmed/26689677

16. Servant G, Weiner OD, Neptune ER, Sedat JW, Bourne HR. Dynamics of a chemoattractant receptor in living neutrophils during chemotaxis. Mol. Biol. Cell [Internet]. 1999;10:1163–78. Available from: http://www.ncbi.nlm.nih.gov/pubmed/10198064

17. Majumdar R, Tavakoli Tameh A, Parent CA. Exosomes Mediate LTB4 Release during Neutrophil Chemotaxis. PLoS Biol. 2016;14:1–28.

18. Yaseen R, Blodkamp S, Lüthje P, Reuner F, Völlger L, Naim HY, et al. Antimicrobial activity of HL-60 cells compared to primary blood-derived neutrophils against Staphylococcus aureus. J. Negat. Results Biomed. [Internet]. 2017;16:2. Available from: https://www.ncbi.nlm.nih.gov/pmc/articles/PMC5316427/

19. Collins SJ. The HL-60 promyelocytic leukemia cell line: proliferation, differentiation, and cellular oncogene expression. Blood. 1987;70:1233–44.

20. Friend C, Scher W, Holland JG, T S. Hemoglobin synthesis in murine virus-induced leukemic cells in vitro: stimulation of erythroid differentiation by dimethyl sulfoxide. Proc. Natl. Acad. Sci. 1971;68:378–82.

21. Siebenlist U, Bressler P, Kelly K. Two distinct mechanisms of transcriptional control operate on c-myc during differentiation of HL60 cells. Mol Cell Biol. 1988;8:867–74.

22. Jorgenson KF, Antoun GR, Zipf TF. Chromatin structural analysis of the 5′ end and contiguous flanking region of the myeloperoxidase gene. Blood. 1991;77:159–64.

23. Veselska R, Zitterbart K, Auer J, Neradil J. Differentiation of HL-60 myeloid leukemia cells induced by all-trans retinoic acid is enhanced in combination with caffeic acid. Int. J. Mol. Med. [Internet]. 2004;14:305–10. Available from: http://www.spandidos-publications.com/10.3892/ijmm.14.2.305

24. Pedruzzi E, Fay M, Elbim C, Gaudry M, Gougerot-Pocidalo M-A. Differentiation of PLB-985 myeloid cells into mature neutrophils, shown by degranulation of terminally differentiated compartments in response to N-formyl peptide and priming of superoxide anion production by granulocyte-macrophage colony-stimulating fact. Br. J. Haematol. [Internet]. 2002;117:719–26. Available from: http://www.ncbi.nlm.nih.gov/pubmed/12028049

25. Nakajima H, Ihle JN. Granulocyte colony-stimulating factor regulates myeloid differentiation through CCAAT/enhancer-binding protein epsilon. Blood [Internet]. 2001;98:897–905. Available from: http://www.ncbi.nlm.nih.gov/pubmed/11493431

26. Collins SR, Yang HW, Bonger KM, Guignet EG, Wandless TJ, Meyer T. Using light to shape chemical gradients for parallel and automated analysis of chemotaxis. Mol. Syst. Biol. [Internet]. 2015;11:804. Available from: http://www.pubmedcentral.nih.gov/articlerender.fcgi?artid=4422560&tool=pmcentrez&rendertype=abstract

27. Marin-Esteban V, Turbica I, Dufour G, Semiramoth N, Gleizes A, Gorges R, et al. Afa/Dr Diffusely Adhering Escherichia coli Strain C1845 Induces Neutrophil Extracellular Traps That Kill Bacteria and Damage Human Enterocyte-Like Cells. Infect. Immun. [Internet]. 2012;80:1891–9. Available from: http://iai.asm.org/content/80/5/1891

28. Chang HH, Oh PY, Ingber DE, Huang S. Multistable and multistep dynamics in neutrophil differentiation. BMC Cell Biol. [Internet]. 2006;7:11. Available from: http://www.ncbi.nlm.nih.gov/pubmed/16507101

29. Le Cabec V, Calafat J, Borregaard N. Sorting Of The Specific Granule Protein, Ngal, During Granulocytic Maturation Of Hl-60 Cells. Blood. 1997;89:2113–21.

30. Meyer PA, Kleinschnitz C. Retinoic Acid Induced Differentiation and Commitment in HL-60 Cells. 1990;88:179–82.

31. Yu L, Zhen L, Dinauer MC. Biosynthesis of the Phagocyte NADPH Oxidase Cytochrome b 558. J. Biol. Chem. 1997;272:27288–94.

32. Bunce CM, French PJ, Durham J, Stockley RA, Michell RH, Brown G. Indomethacin potentiates the induction of HL60 differentiation to neutrophils, by retinoic acid and granulocyte colony-stimulating factor, and to monocytes, by vitamin D3. Leukemia. 1994;8:595–604.

33. Fleck RA, Athwal H, Bygraves JA, Hockley DJ, Feavers IM, Stacey GN. Optimization of nb-4 and hl-60 differentiation for use in opsonophagocytosis assays. In Vitro Cell. Dev. Biol. Anim. 2003;39:235–42.

34. Huang S, Eichler G, Bar-Yam Y, Ingber DE. Cell fates as high-dimensional attractor states of a complex gene regulatory network. Phys. Rev. Lett. 2005;94:1–4.

35. Jiang K, Zhu L, Buck MJ, Chen Y, Carrier B, Liu T, et al. Disease-Associated Single-Nucleotide Polymorphisms From Noncoding Regions in Juvenile Idiopathic Arthritis Are Located Within or Adjacent to Functional Genomic Elements of Human Neutrophils and CD4+ T Cells. Arthritis Rheumatol. [Internet]. 2015;67:1966–77. Available from: http://www.ncbi.nlm.nih.gov/pubmed/25833190

36. Wright HL, Thomas HB, Moots RJ, Edwards SW. RNA-seq reveals activation of both common and cytokine-specific pathways following neutrophil priming. PLoS One [Internet]. 2013;8:e58598. Available from: http://www.pubmedcentral.nih.gov/articlerender.fcgi?artid=3590155&tool=pmcentrez&rendertype=abstract

37. Thomas HB, Moots RJ, Edwards SW, Wright HL. Whose gene is it anyway? the effect of preparation purity on neutrophil transcriptome studies. PLoS One. 2015;10:1–15.

38. Cusanovich DA, Daza R, Adey A, Pliner HA, Christiansen L, Gunderson KL, et al. Multiplex single-cell profiling of chromatin accessibility by combinatorial cellular indexing. Science [Internet]. 2015;348:910–4. Available from: http://www.ncbi.nlm.nih.gov/pubmed/25953818

39. Taylor A, Tang W, Bruscia EM, Zhang PX, Lin A, Gaines P, et al. SRF is required for neutrophil migration in response to inflammation. Blood. 2014;123:3027–36.

40. Coffelt SB, Kersten K, Doornebal CW, Weiden J, Vrijland K, Hau C-S, et al. IL-17-producing y6 T cells and neutrophils conspire to promote breast cancer metastasis. Nature. 2015;522:345–8.

41. NCBI Resource Coordinators. Database Resources of the National Center for Biotechnology Information. Nucleic Acids Res. [Internet]. 2017;45:D12–7. Available from: http://www.ncbi.nlm.nih.gov/pubmed/27899561

42. Huang DW, Sherman BT, Lempicki RA. Systematic and integrative analysis of large gene lists using DAVID bioinformatics resources. Nat. Protoc. [Internet]. 2008 [cited 2017 Sep 18];4:44–57. Available from: http://www.ncbi.nlm.nih.gov/pubmed/19131956

43. Van der Auwera GA, Carneiro MO, Hartl C, Poplin R, del Angel G, Levy-Moonshine A, et al. From FastQ Data to High-Confidence Variant Calls: The Genome Analysis Toolkit Best Practices Pipeline. Curr. Protoc. Bioinforma. [Internet]. Hoboken, NJ, USA: John Wiley & Sons, Inc.; 2013. p. 11.10.1–11.10.33. Available from: http://www.ncbi.nlm.nih.gov/pubmed/25431634

44. Dobin A, Davis CA, Schlesinger F, Drenkow J, Zaleski C, Jha S, et al. STAR: Ultrafast universal RNA-seq aligner. Bioinformatics. 2013;29:15–21.

45. Trapnell C, Williams BA, Pertea G, Mortazavi A, Kwan G, van Baren MJ, et al. Transcript assembly and quantification by RNA-Seq reveals unannotated transcripts and isoform switching during cell differentiation. Nat. Biotechnol. Nature Publishing Group; 2010;28:511–5.

46. Glusman G, Caballero J, Robinson M, Kutlu B, Hood L. Optimal Scaling of Digital Transcriptomes. PLoS One [Internet]. 2013;8:e77885. Available from: http://dx.plos.org/10.1371/journal.pone.0077885

